## Supplemental Figures and Tables for "First report and characterization of a plasmid-encoded *bla*_SFO-1_ in a multi-drug resistant *Aeromonas hydrophila* clinical isolate"

### Supplemental Table 1. Genes used to perform phylogenetic analysis of MAH-4 with representative *Aeromonas* species

| Product | PGFam | Align. Score | Align. Length | Num Seqs | Mean Sqr Freq | Prop Gaps |
|---|---|---|---|---|---|---|
| Glycine dehydrogenase [decarboxylating] (glycine cleavage system P protein) (EC 1.4.4.2) | PGF_00008773 | 29.08 | 958 | 26 | 0.94 | 0.007 |
| DNA gyrase subunit B (EC 5.99.1.3) | PGF_06703483 | 27.57 | 804 | 26 | 0.972 | 0.001 |
| BioD-like N-terminal domain / Phosphate acetyltransferase (EC 2.3.1.8) | PGF_01937476 | 25.61 | 727 | 26 | 0.95 | 0.02 |
| Oligopeptidase A (EC 3.4.24.70) | PGF_00026904 | 23.9 | 680 | 26 | 0.917 | 0.011 |
| Sensory box sensor/GGDEF/EAL domain protein | PGF_00050892 | 23.37 | 893 | 26 | 0.782 | 0.072 |
| DNA primase DnaG | PGF_00421679 | 23.1 | 596 | 26 | 0.946 | 0.002 |
| Phosphoenolpyruvate carboxykinase [ATP] (EC 4.1.1.49) | PGF_02947204 | 22.28 | 546 | 26 | 0.954 | 0.009 |
| NAD synthetase (EC 6.3.1.5) / Glutamine amidotransferase chain of NAD synthetase | PGF_06525160 | 22.05 | 540 | 26 | 0.949 | 0 |
| Alkyl hydroperoxide reductase protein F | PGF_09969389 | 22.04 | 530 | 26 | 0.958 | 0.006 |
| Hydroxylamine reductase (EC 1.7.99.1) | PGF_00736089 | 21.88 | 550 | 26 | 0.933 | 0.002 |
| 2-isopropylmalate synthase (EC 2.3.3.13) | PGF_04294677 | 21.88 | 524 | 26 | 0.956 | 0.004 |
| Guanosine-5'-triphosphate,3'-diphosphate pyrophosphatase (EC 3.6.1.40) @ Exopolyphosphatase (EC 3.6.1.11) | PGF_05837876 | 21.26 | 498 | 26 | 0.952 | 0.004 |
| ATP synthase beta chain (EC 3.6.3.14) | PGF_05195027 | 21.1 | 462 | 26 | 0.982 | 0 |
| Methylcrotonyl-CoA carboxylase carboxyl transferase subunit (EC 6.4.1.4) | PGF_05369673 | 20.91 | 549 | 26 | 0.892 | 0.025 |
| IMP cyclohydrolase (EC 3.5.4.10) / Phosphoribosylaminoimidazolecarboxamide formyltransferase (EC 2.1.2.3) | PGF_00013509 | 20.85 | 535 | 26 | 0.901 | 0.033 |
| Glutamate-1-semialdehyde 2,1-aminomutase (EC 5.4.3.8) | PGF_09019918 | 19.97 | 429 | 26 | 0.964 | 0.002 |
| tRNA cytosine(34) acetyltransferase (EC 2.3.1.193) | PGF_01956528 | 19.95 | 736 | 26 | 0.735 | 0.065 |
| O-acetylhomoserine sulfhydrylase (EC 2.5.1.49) @ O-succinylhomoserine sulfhydrylase (EC 2.5.1.48) | PGF_04792526 | 19.83 | 422 | 26 | 0.965 | 0 |
| Paraquat-inducible protein B | PGF_12719092 | 19.81 | 549 | 26 | 0.845 | 0.022 |
| ATP-dependent RNA helicase RhlB (EC 3.6.4.13) | PGF_02213711 | 19.75 | 429 | 26 | 0.954 | 0.011 |
| Outer membrane stress sensor protease DegQ, serine protease | PGF_03779043 | 19.67 | 469 | 26 | 0.908 | 0.031 |
| Aromatic amino acid transport protein AroP | PGF_00064863 | 19.64 | 460 | 26 | 0.916 | 0.007 |
| Putative Na+/H+ antiporter | PGF_10543494 | 19.25 | 444 | 26 | 0.914 | 0.037 |
| Adenylosuccinate synthetase (EC 6.3.4.4) | PGF_06935032 | 19.21 | 418 | 26 | 0.939 | 0.002 |
| Membrane-bound lytic murein transglycosylase F (EC 4.2.2.n1) | PGF_00020190 | 19.12 | 508 | 26 | 0.848 | 0.028 |
| Murein peptide ABC transporter, substrate-binding protein (requires DppBCDF) | PGF_00030901 | 18.93 | 553 | 26 | 0.805 | 0.056 |
| Murein DD-endopeptidase MepM | PGF_00023403 | 18.71 | 448 | 26 | 0.884 | 0.013 |

| | | | | | | |
|---|---|---|---|---|---|---|
| Two-component system sensor histidine kinase | PGF_00052313 | 18.7 | 449 | 26 | 0.882 | 0.031 |
| Sodium/glutamate symporter | PGF_06816941 | 18.58 | 440 | 26 | 0.886 | 0.029 |
| Probable low-affinity inorganic phosphate transporter | PGF_00037171 | 18.56 | 421 | 26 | 0.905 | 0.026 |
| GTP-binding and nucleic acid-binding protein YchF | PGF_00007012 | 18.46 | 363 | 26 | 0.969 | 0 |
| Succinyl-CoA ligase [ADP-forming] beta chain (EC 6.2.1.5) | PGF_00054528 | 18.27 | 388 | 26 | 0.927 | 0.025 |
| Lipoprotein releasing system transmembrane protein LolC | PGF_08074580 | 17.93 | 439 | 26 | 0.856 | 0.07 |
| Integral membrane protein | PGF_00557891 | 17.91 | 393 | 26 | 0.903 | 0.016 |
| Probable Na+/H+-dicarboxylate symporter | PGF_00552628 | 17.88 | 399 | 26 | 0.895 | 0.026 |
| 6-phosphofructokinase (EC 2.7.1.11) | PGF_07421229 | 17.45 | 323 | 26 | 0.971 | 0 |
| Biotin synthase (EC 2.8.1.6) | PGF_01400330 | 17.33 | 392 | 26 | 0.875 | 0.075 |
| Glycyl-tRNA synthetase alpha chain (EC 6.1.1.14) | PGF_00009967 | 17.3 | 310 | 26 | 0.983 | 0.009 |
| Porphobilinogen synthase (EC 4.2.1.24) | PGF_00489714 | 17.3 | 339 | 26 | 0.939 | 0.001 |
| Transcriptional factor MdcH | PGF_00559497 | 17.22 | 333 | 26 | 0.944 | 0.014 |
| Cytochrome c heme lyase subunit CcmH | PGF_06068352 | 17.16 | 427 | 26 | 0.831 | 0.012 |
| Deoxyguanosinetriphosphate triphosphohydrolase (EC 3.1.5.1), subgroup 1 | PGF_10089955 | 17.15 | 458 | 26 | 0.801 | 0.046 |
| Phosphate ABC transporter, substrate-binding protein PstS (TC 3.A.1.7.1) | PGF_07668761 | 17.07 | 324 | 26 | 0.948 | 0 |
| Branched-chain amino acid ABC transporter, permease protein LivH (TC 3.A.1.4.1) | PGF_06868199 | 17.03 | 308 | 26 | 0.97 | 0 |
| Cell-division-associated, ABC-transporter-like signaling protein FtsX | PGF_03701810 | 16.96 | 318 | 26 | 0.951 | 0.003 |
| Flagellar protein FlgT | PGF_09781823 | 16.8 | 431 | 26 | 0.809 | 0.095 |
| Transcriptional activator NhaR | PGF_00057513 | 16.58 | 310 | 26 | 0.942 | 0.014 |
| Holliday junction ATP-dependent DNA helicase RuvB (EC 3.6.4.12) | PGF_05049118 | 16.58 | 337 | 26 | 0.903 | 0.031 |
| Glycerol-3-phosphate dehydrogenase [NAD(P)+] (EC 1.1.1.94) | PGF_00008611 | 16.41 | 334 | 26 | 0.898 | 0.035 |
| Tryptophanase (EC 4.1.99.1) | PGF_03720595 | 16.38 | 490 | 26 | 0.74 | 0.041 |
| tRNA dimethylallyltransferase (EC 2.5.1.75) | PGF_04807486 | 16.24 | 309 | 26 | 0.924 | 0.002 |
| (2E,6E)-farnesyl diphosphate synthase (EC 2.5.1.10) | PGF_06784545 | 16.06 | 296 | 26 | 0.934 | 0 |
| Formyltetrahydrofolate deformylase (EC 3.5.1.10) | PGF_00006320 | 15.94 | 283 | 26 | 0.948 | 0.015 |
| 33 kDa chaperonin HslO | PGF_10462808 | 15.88 | 295 | 26 | 0.925 | 0.003 |
| Lipoprotein NlpI | PGF_00017407 | 15.76 | 319 | 26 | 0.883 | 0.037 |
| Cardiolipin synthase (EC 2.7.8.-) phosphatidylethanolamine-utilizing, bacterial type ClsC | PGF_08028591 | 15.67 | 437 | 26 | 0.75 | 0.044 |
| tRNA (guanine(37)-N(1))-methyltransferase (EC 2.1.1.228) | PGF_00413208 | 15.57 | 249 | 26 | 0.987 | 0.001 |
| Uncharacterized efflux ABC transporter, ATP-binding protein YadG | PGF_06025565 | 15.47 | 334 | 26 | 0.847 | 0.09 |
| Outer membrane porin OmpC | PGF_00027880 | 15.42 | 412 | 26 | 0.76 | 0.137 |

| | | | | | | |
|---|---|---|---|---|---|---|
| Phospholipid ABC transporter permease protein MlaE | PGF_07114837 | 15.38 | 259 | 26 | 0.956 | 0.004 |
| FIG008480: zinc-binding hydrolase | PGF_02211096 | 15.27 | 265 | 26 | 0.938 | 0 |
| Uridylate kinase (EC 2.7.4.22) | PGF_02923127 | 15.15 | 244 | 26 | 0.97 | 0.002 |
| Rod shape-determining protein MreC | PGF_10369954 | 15.09 | 312 | 26 | 0.854 | 0.069 |
| Bis(5'-nucleosyl)-tetraphosphatase, symmetrical (EC 3.6.1.41) | PGF_07763915 | 15.08 | 275 | 26 | 0.909 | 0.007 |
| Cytochrome c-type biogenesis protein CcmC, putative heme lyase for CcmE | PGF_00420482 | 14.99 | 246 | 26 | 0.956 | 0 |
| tRNA uridine 5-oxyacetic acid(34) methyltransferase (EC 2.1.1.-) | PGF_00049344 | 14.93 | 271 | 26 | 0.907 | 0.04 |
| Formamidopyrimidine-DNA glycosylase (EC 3.2.2.23) | PGF_05740384 | 14.91 | 270 | 26 | 0.907 | 0.023 |
| Tol-Pal system protein TolQ | PGF_00648054 | 14.75 | 229 | 26 | 0.975 | 0.001 |
| Arginine ABC transporter, substrate-binding protein ArtI | PGF_00502921 | 14.71 | 246 | 26 | 0.938 | 0.004 |
| Ribose ABC transporter, substrate-binding protein RbsB (TC 3.A.1.2.1) | PGF_12669939 | 14.64 | 356 | 26 | 0.776 | 0.181 |
| Pyrroline-5-carboxylate reductase (EC 1.5.1.2) | PGF_07609122 | 14.62 | 276 | 26 | 0.88 | 0.007 |
| Peptide ABC transporter, ATP-binding protein SapF | PGF_08049323 | 14.59 | 279 | 26 | 0.874 | 0.064 |
| Histidine ABC transporter, permease protein HisM (TC 3.A.1.3.1) | PGF_00974916 | 14.43 | 230 | 26 | 0.952 | 0 |
| hypothetical protein | PGF_00195010 | 14.43 | 287 | 26 | 0.852 | 0.007 |
| NADPH-dependent 7-cyano-7-deazaguanine reductase (EC 1.7.1.13) | PGF_00024855 | 14.39 | 302 | 26 | 0.828 | 0.061 |
| Zn-dependent protease with chaperone function PA4632 | PGF_01922821 | 14.26 | 277 | 26 | 0.857 | 0.022 |
| Ribonuclease III (EC 3.1.26.3) | PGF_03790040 | 14.25 | 224 | 26 | 0.952 | 0.005 |
| Uncharacterized UPF0721 integral membrane protein | PGF_08139483 | 14.18 | 262 | 26 | 0.876 | 0.019 |
| Stringent starvation protein A | PGF_00054301 | 14.13 | 209 | 26 | 0.977 | 0 |
| 23S rRNA (adenine(1618)-N(6))-methyltransferase (EC 2.1.1.181) | PGF_00415130 | 13.98 | 338 | 26 | 0.761 | 0.067 |
| Oxidoreductase, Gfo/Idh/MocA family | PGF_00738912 | 13.91 | 313 | 26 | 0.786 | 0.037 |
| Arginyl-tRNA--protein transferase (EC 2.3.2.8) | PGF_02992100 | 13.88 | 238 | 26 | 0.9 | 0.005 |
| FIG000875: Thioredoxin domain-containing protein EC-YbbN | PGF_03863088 | 13.72 | 282 | 26 | 0.817 | 0.024 |
| NADH pyrophosphatase (EC 3.6.1.22), decaps 5'-NAD modified RNA | PGF_00024615 | 13.63 | 267 | 26 | 0.834 | 0.03 |
| FIG00003370: Multicopper polyphenol oxidase | PGF_05501316 | 13.61 | 251 | 26 | 0.859 | 0.022 |
| Inner membrane protein YohK | PGF_00014196 | 13.46 | 234 | 26 | 0.88 | 0.026 |
| Predicted hydrolase/acyltransferase | PGF_03831039 | 13.43 | 292 | 26 | 0.786 | 0.021 |
| Outer membrane beta-barrel assembly protein BamD | PGF_05872952 | 13.43 | 305 | 26 | 0.769 | 0.164 |
| rRNA (guanine-N(1))-methyltransferase | PGF_00556937 | 13.37 | 292 | 26 | 0.782 | 0.086 |
| Translation elongation factor P | PGF_03799365 | 13.33 | 188 | 26 | 0.972 | 0 |
| Phospholipid ABC transporter shuttle protein MlaC | PGF_10659937 | 13.33 | 211 | 26 | 0.918 | 0.005 |

| | | | | | | |
|---|---|---|---|---|---|---|
| LSU ribosomal protein L5p (L11e) | PGF_00016443 | 13.26 | 179 | 26 | 0.991 | 0 |
| Polyphosphate glucokinase (EC 2.7.1.63) | PGF_04211832 | 13.25 | 265 | 26 | 0.814 | 0.076 |
| Outer membrane protein OmpK | PGF_00028291 | 13.09 | 294 | 26 | 0.764 | 0.057 |
| Quercetin 2,3-dioxygenase (EC 1.13.11.24) => YhhW | PGF_02380724 | 13.08 | 228 | 26 | 0.867 | 0.025 |
| UDP-2,3-diacylglucosamine diphosphatase (EC 3.6.1.54) | PGF_00063937 | 13.08 | 254 | 26 | 0.821 | 0.048 |
| Outer membrane porin OmpC | PGF_00557072 | 12.91 | 419 | 26 | 0.631 | 0.16 |
| hypothetical protein | PGF_00255967 | 12.7 | 264 | 26 | 0.782 | 0.044 |
| Putative transmembrane protein | PGF_01769544 | 12.69 | 186 | 26 | 0.931 | 0.007 |

### Supplemental Table 2. Nucleotide sequence similarity to *A. hydrophila* pMAH-4 using BLAST®

| Description | max score | Total score | Query cover | E-value | % identity | Accession length | Acc. # |
|---|---|---|---|---|---|---|---|
| Aeromonas caviae strain AC1520 plasmid pAC1520, complete sequence | 66550 | 4.54E+05 | 75% | 0 | 95.88% | 253471 | CP120943.1 |
| Klebsiella pneumoniae strain KP1814 plasmid pKP1814-1, complete sequence | 48230 | 1.35E+05 | 20% | 0 | 100.00% | 299858 | KX839207.1 |
| Klebsiella variicola strain 4253 plasmid p4253-imp, complete sequence | 48230 | 1.36E+05 | 19% | 0 | 100.00% | 334271 | CP135069.1 |
| Klebsiella variicola strain SHET-01 plasmid pNDM-IMP-1, complete sequence | 48230 | 1.45E+05 | 20% | 0 | 100.00% | 347317 | CP050681.1 |
| Klebsiella pneumoniae strain WH11 plasmid pWH11, complete sequence | 48230 | 1.03E+05 | 16% | 0 | 100.00% | 325030 | ON882017.1 |
| Enterobacter asburiae strain AR2284-yvys plasmid pAR2284_1, complete sequence | 48230 | 1.62E+05 | 21% | 0 | 100.00% | 373545 | CP083831.1 |
| Klebsiella quasipneumoniae strain SWMUF35 plasmid pA, complete sequence | 48230 | 1.35E+05 | 20% | 0 | 100.00% | 311723 | CP068445.1 |
| Klebsiella pneumoniae strain KP294 plasmid pIMP4-KP294, complete sequence | 48224 | 1.53E+05 | 20% | 0 | 99.99% | 349403 | CP083446.1 |
| Klebsiella pneumoniae strain KP19-2581 plasmid pKP19-2581_367k_HI5_, complete sequence | 45493 | 85100 | 9% | 0 | 100.00% | 367802 | CP120875.1 |
| Citrobacter freundii strain IDR1800045912-01-00 plasmid p1C157, complete sequence | 39506 | 66744 | 9% | 0 | 100.00% | 156725 | CP054295.1 |
| Aeromonas caviae strain SCLZS52 chromosome, complete genome | 39500 | 1.74E+05 | 21% | 0 | 99.99% | 4718963 | CP091176.1 |
| Klebsiella pneumoniae plasmid pQL5-NDM-KPC, complete sequence | 36863 | 81155 | 7% | 0 | 99.99% | 214266 | OR253888.1 |
| Pseudomonas mandelii JR-1 plasmid, complete sequence | 29288 | 40648 | 6% | 0 | 99.99% | 410512 | CP005961.1 |
| Aeromonas caviae DNA, complete genome, strain: WP3-S18-ESBL-02 | 28987 | 1.08E+05 | 14% | 0 | 99.94% | 4860108 | AP022013.1 |
| Klebsiella quasipneumoniae strain C3001 plasmid pC3001-2-NDM, complete sequence | 28509 | 33544 | 5% | 0 | 100.00% | 270400 | CP039792.1 |
| Klebsiella quasipneumoniae strain A708 plasmid pA708-1, complete sequence | 28507 | 81589 | 12% | 0 | 100.00% | 238703 | CP026369.1 |
| Klebsiella aerogenes strain AR_0161 plasmid unnamed, complete sequence | 28507 | 1.07E+05 | 15% | 0 | 100.00% | 451422 | CP028952.1 |
| Klebsiella pneumoniae strain A708 plasmid pA708-IMP, complete sequence | 28507 | 81589 | 12% | 0 | 100.00% | 238703 | MF344567.1 |
| Klebsiella quasipneumoniae subsp. quasipneumoniae plasmid p2019SCSN059_tmexCD_333k, complete sequence | 28502 | 1.19E+05 | 15% | 0 | 99.99% | 333095 | ON169978.1 |
| Klebsiella pneumoniae strain KP21300 plasmid pIMP_KP21300, complete sequence | 28498 | 85728 | 11% | 0 | 99.98% | 279261 | CP124694.1 |
| Klebsiella variicola plasmid pFK2020ZBJ35_tmexCD_325k, complete sequence | 27835 | 55700 | 9% | 0 | 99.99% | 325393 | ON169979.1 |
| Klebsiella michiganensis strain 7525 plasmid pKOX7525_1, complete sequence | 27215 | 1.33E+05 | 18% | 0 | 99.99% | 397447 | CP065475.1 |
| Aeromonas sp. ASNIH2 chromosome, complete genome | 26328 | 86766 | 13% | 0 | 99.99% | 4801408 | CP026406.1 |
| Escherichia coli strain 1585m1 plasmid p1585m1_B, complete sequence | 25481 | 47157 | 6% | 0 | 99.96% | 113047 | CP086388.1 |
| Citrobacter freundii strain Cf.1 chromosome, complete genome | 25481 | 39440 | 6% | 0 | 99.96% | 5153556 | CP085642.1 |
| Klebsiella pneumoniae strain QD23 chromosome, complete genome | 25475 | 1.75E+05 | 9% | 0 | 99.96% | 5803733 | CP042858.1 |
| Citrobacter freundii strain Cf52 plasmid pCf52, complete sequence | 25475 | 77541 | 8% | 0 | 99.96% | 219342 | KY887592.1 |
| Citrobacter freundii strain F3517 plasmid pF3517-1, complete sequence | 25475 | 54392 | 7% | 0 | 99.96% | 139338 | CP137177.1 |
| Citrobacter freundii strain Survcare457 plasmid pCF_Surv457-KPC2, complete sequence | 25473 | 44814 | 6% | 0 | 99.96% | 83074 | CP104961.1 |
| Klebsiella pneumoniae strain KP1572 plasmid pIMP1572, complete sequence | 25470 | 65626 | 7% | 0 | 99.95% | 142993 | MH464586.1 |
| Enterobacter hormaechei strain E5 plasmid pE5_003, complete sequence | 25470 | 40243 | 5% | 0 | 99.95% | 86404 | CP042574.1 |
| Aeromonas hydrophila strain K522 plasmid pK522-MOX, complete sequence | 25470 | 43627 | 7% | 0 | 99.95% | 176681 | CP118701.1 |

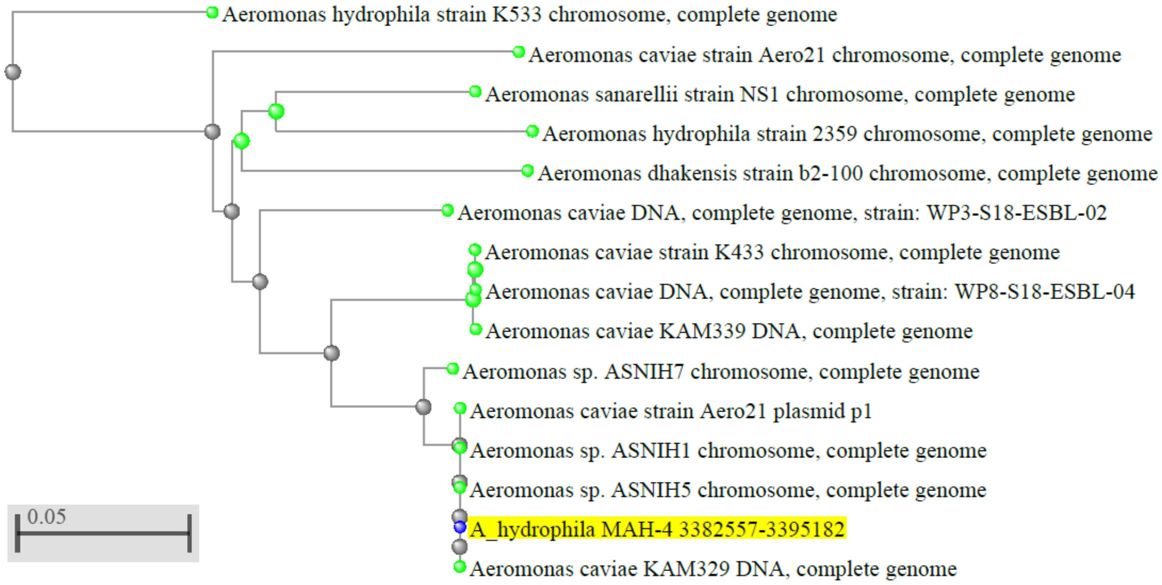

Supplemental Figure 1. Phylogenetic analysis of Trb operon. BLAST Tree was used to compared the chromosomal DNA encoding a Trb operon (trbK-virD4-trbBCDEJKLFGI: 3,382,557-3,395,182) from *Aeromonas hydrophila* MAH-4 against members of the Aeromonadaceae/Succinivibrionaceae group (taxid:135,624).

Table S3. Mobile genetic elements on *Aeromonas hydrophila* pMAH-1

| MGE | MGE family | Location | Origin | Acc.# | Score (bits) | E-value |
|---|---|---|---|---|---|---|
| ISPst3 | IS21 | 325,934 - 992 | *Pseudomonas stutzeri* | AB088753 | 4726 | 0.0 |
| ISPa60 | ISAs1 | 1,072 – 2,280 | *Pseudomonas aeruginosa* | | 2381 | 0.0 |
| ISAhy2 | IS630 | 2,543 – 3,700 | *Aeromonas hydrophila* | FM877486 | 2256 | 0.0 |
| IS6100 | IS6 | 30,878 – 31,703 | *Mycobacterium fortuitum* | X53635 | 1637 | 0.0 |
| IS26 | IS6 | 34,991 – 35,810 | *Proteus vulgaris* | X00011 | 1624 | 0.0 |
| ISCfr1 | IS1182 | 39,539 – 41,155 | *Citrobacter freundii* | AF550415 | 3205 | 0.0 |
| IS6100 | IS6 | 52,786 – 53,665 | *Mycobacterium fortuitum* | X53635 | 1744 | 0.0 |
| TnAs1 | Tn3 | 42,984 - 45,884 | *Aeromonas salmoncida* | CP022426 | 5618 | 0.0 |
| Tn6082 | Tn3 | 15,120 – 18,005 | *Erwinia amylovora* | M96392 | 5642 | 0.0 |
| **Composite transposons** | | | | | | |
| IS6100 - flanked | | 30,877 - 53,665 | | | | |
